# MAGICTRICKS: A tool for predicting transcription factors and cofactors that drive gene lists

**DOI:** 10.1101/492744

**Authors:** A Roopra

## Abstract

Transcriptomic profiling is an immensely powerful hypothesis generating tool. However, accurately predicting the transcription factors (TFs) and cofactors that drive transcriptomic differences between samples represents a challenge and current approaches are limited by high false discovery rates. This is due to the use of TF binding sequence motifs that, due their small size, are found randomly throughout the genome, and do not allow discovery of cofactors. A second limitation is that even the most advanced approaches that use ChIPseq tracks hosted at sites such as the Encyclopedia Of DNA Elements (ENCODE) assign TFs and cofactors to genes via a binary designation of ‘target’, or ‘non-target’ that ignores the intricacies of the biology behind transcriptional regulation.

ENCODE archives ChIPseq tracks of 169 TFs and cofactors assayed in 91 cell lines. The algorithm presented herein, **M**ining **G**ene **C**ohorts for **T**ranscriptional **R**egulators **I**nferred by **K**olmogorov-Smirnov **S**tatistics (MAGICTRICKS), uses ENCODE ChIPseq data to look for statistical enrichment of TFs and cofactors in gene bodies and flanking regions in gene lists. When compared to 2 commonly used web resources, o-Possum and Enrichr, MAGICTRICKS was able to more accurately predict TFs and cofactors that drive gene changes in 3 settings: 1) A cell line expressing or lacking single TF, 2) Breast tumors divided along PAM50 designations and 3) Whole brain samples from WT mice or mice lacking a single TF in a particular neuronal subtype. In summary, MAGICTRICKS is a standalone application that runs on OSX and Windows machines that produces meaningful predictions of which TFs and cofactors are enriched in a gene list.

## Introduction

Key to the control of gene expression is the level of transcript in the cell. Transcription factors (TFs) are DNA binding proteins that recognize specific sequence elements (motifs) associated with gene promoters and enhancers. TFs recruit cofactors that do not themselves bind DNA but are brought to promoters via TFs to either enhance or repress gene expression. TFs and cofactors (here on termed ‘Factors’) are thus key regulators of transcript levels.

It is now routine to obtain the levels of every gene transcript in the genome i.e. whole transcriptome data. The datasets contain tens of thousands of expression values per sample and the number of samples can be in the thousands such as breast cancer transcriptomes archived at The Cancer Genome Atlas (TCGA)^1^. When comparing transcriptomes from two or more conditions such as normal to cancerous tissue, thousands of mRNA levels can change. We posit that in many cases, the majority of those changes are driven by alterations in the function of a few Factors that coordinate programmatic gene changes on a genome wide scale. Identifying those driving Factors is a fundamental problem and therapeutic opportunity, yet current tools are extremely poor at making such predictions.

Current algorithms fall into two categories. The first looks for motifs that are over-represented in the upstream regions of a list of genes with altered expression compared to the gene population. This approach often fails because with very few exceptions, transcription factor motifs are small (around 6bp) and often have redundant bases. A random 6bp sequence occurs approximately 750,000 times in the human genome. Thus, searches will yield a high false positive rate; most predicted sites will not be functional. Moreover, in many genes, TFs act in concert such that the presence of a single motif element is uninformative. Motif searching also precludes any prediction of cofactors because cofactors do not bind DNA directly. This is especially problematic for translational research where uncovering drugable targets is a major goal, and cofactors are often enzymes that can make excellent drug targets. The second approach relies on assigning genes to Factors (TFs and cofactors) by looking at ChIPSeq tracks such as those archived at ENCODE^1^. Genes are ranked by ChIP signal and then an arbitrary number of ‘top’ genes are taken as targets. This approach is exemplified by Enrichr^2^ Enrichr is a very powerful tool with excellent performance that allows for assignment of cofactors. However, in general, the binary assignment of genes as targets or non-targets of a Factor does not take into account the biology of Factor regulation of genes. Many Factors have highly non-gaussian ChIP signal distributions; the distributions can have large numbers of sites with very low but detectable signals and skewed, long tails of decreasing ChIP signal. This thwarts unbiased attempts to define genes as ‘targets’ or ‘non-targets’^3^. A new method is required that takes into account the biology of transcriptional regulation in a statistically rigorous manner.

We have devised a novel algorithm that meshes whole transcriptome data with whole genome factor binding (ChIP Seq) data archived at ENCODE. The algorithm is termed ***M**ining **G**ene **C**ohorts for **T**ranscriptional **R**egulators **I**nferred by **K**olmogorov-Smirnov **S**tatistics* (MAGICTRICKS). MAGICTRICKS circumvents the principal confounds of current methods to identify Factors, namely: 1) unacceptably high false positive rates due to the use of TF motif searches 2) inability to identify cofactors due the absence of any binding site motifs 3) arbitrary assignment of Factors to genes based on hard cutoffs of ChIPSeq signals. MAGICTRICKS accepts an input list of genes and compares ChIP signals for the list with the population of ChIP signals for each Factor in ENCODE. We compared MAGICTRICKS to 2 commonly used TF prediction alorithms, oPOSSUM-3^4^ and Enrichr^2^ using whole transcriptome data from a REST knockdown cell line, TCGA breast tumors and mouse brain tissue. In all cases, MAGICTRICKS predicted biologically meaningful TFs and associated cofactors.

## Methods

### MCF7 RNA profiling

MCF7 cells harboring a REST shRNA knockdown and controls were generated as described^5^. RNA was extracted in triplicate from shControl and shREST cells using TRIZOL, checked on an Agilent NanoChip and subjected to microarray hybridization onto Nimblegen arrays (2006-08-03_HG18_60mer_expr) at the University of Wisconsin-Madison Biotechnology Center. Spot intensities were RMA normalized^6^. Probes were considered present in the experiment if they were ‘present’ in all 3 samples of either controls or knockdown cells. If a probe was present in only one condition, the values in the other condition were accepted regardless of the present/absent call. Probes were then collapsed to a single gene value per sample by taking the median value of all probes per gene per sample. Fold changes between control and REST knock-down cells were calculated along with Benjamini-Hochberg corrected Student t-test p values. ‘Up-regulated’ and ‘down-regulated’ gene lists for MAGICTRICKS analysis were generated by taking any genes changed more than 3 Standard Deviations from the mean fold change and having a corrected p value<0.05.

### TCGA profiling

Breast cancer RNAseq data (data_RNA_Seq_v2_expression_median.txt) was downloaded from cBioPortal (latest datafreeze as of 6July2017). PAM50^7^ designations were used to parse tumors into Luminal-A or Basal subtypes (supplemental File 9 - list of samples and PAM50 designation). A gene with an RPKM ≥1 in at least 50% of either basal or Luminal-A samples was considered for further analysis. Fold changes, statistics and gene lists were then generated as above for REST knock-down cells.

### Sams et al profiling

RNAseq data (GSE84175) was downloaded from GEO. A gene with an RPKM ≥1 in all controls or all CTCF KO cells was considered for further analysis and fold changes, statistics and gene lists were then generated as above for REST knock-down cells.

### MAGICTRICKS

MAGICTRICKS aims to determine whether genes in a query list are associated with higher ChIP values than expected by chance for a given transcription factor or cofactor based on ChIPseq tracks archived at ENCODE. To do this, we assigned a ChIP value for each factor in ENCODE at every gene. We then generated an algorithm to test the hypothesis that a list of query genes are enriched for high ChIP signals.

### Generation of the MAGIC Matrix

We defined a single ChIP signal for each factor in ENCODE at every gene as follows:

All genes in the human genome (hg19) were assigned a gene domain that was defined as 5Kb either side of the gene body (5’UTR − 5Kb to 3’UTR + 5Kb). NarrowPeak files for all transcription factors and cofactors were downloaded from http://genome.ucsc.edu/cgi-bin/hgFileUi?db=hg19&g=wgEncodeAwgTfbsUniform. These files provide called peaks of signal enrichment based on pooled, normalized (interpreted) data. Custom Python scripts were then used to extract the highest ChIPseq peak value (signalValue) for a given factor seen in any cell line within every gene domain. Thus each factor and gene were assigned a single signalValue – the highest value – observed in ENCODE. The highest value was chosen to 1) maximize the likelihood that this site will be bound in cell types not present in ENCODE and 2) Work by others suggest that the strongest sites may be the most functional ^3^. The resulting array (factor columns vs gene rows) was normalized such that the sum of all signalValues for any factor (columns) was arbitrarily set at 100,000. We term the final array the MAGIC Matrix (Supplemental File 1).

### Defining Enriched Factors in a Gene list

Two lists are entered into MAGICTRICKS. The first is a master list containing all genes of relevance to the experiment (for example, all genes deemed to be present or expressed in an experiment). The second list contains genes of interest – the query list (e.g. genes induced above a threshold in one condtion over control). Query lists are subsets of the master list and both lists are filtered by MAGICTRICKS for genes present in the MAGIC matrix.

For each Factor in the Matrix, MAGICTRICKS orders the N genes in the accepted master list by ChIP signal (lowest to highest) obtained from MAGIC Matrix. It then generates a normalized cumulative distribution function (CDF) such that:

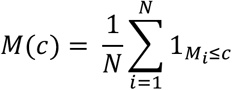

Where M(c) is the fraction of genes in the Master list (N genes) with ChIP signal less than c, and M are the ordered ChIP signals. 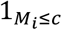 is an indicator function that is equal to 1 if M_i_ ≤ c else equal to 0. An emperical distribution function, Q(c) is generated by similarly ordering the X query genes Q, where Q⊂M, and 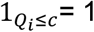 if Q_i_ ≤ c else 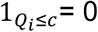:

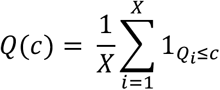

It then screens for Factors where the query distribution is right-shifted compared to the master CDF. This is accomplished by calculating the difference between the 2 distributions and defining the supremum (D_s_) and infimum D_I_ of the difference as:

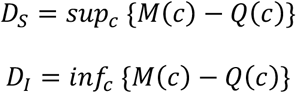

If the cumulative of query samples is left shifted compared to the population (|D_S_| < |D_I_|), the factor is triaged and not considered further. If the query cumulative is right shifted compared to the population cumulative i.e. |D_S_| > |D_I_|, MAGICTRICKS performs a 1-tailed Kolmorogov-Smirnov (KS) test between the master and query list to test the null hypothesis that the query list is drawn randomly from the master; D_S_ equals the KS statistic. The argument of the KS statistic is the ‘critical ChIP’ and represents the minimum ChIP value a gene must have to be considered a target gene for the Factor.

A Score for each Factor is calculated to incorporate the Benjamini-Hochberg corrected KS p value, as well as a measure of how the highest ChIP values in the query list compare to the highest values in the population. Thus, for each Factor, the mean of the top n chip signals in the query list is compared to the mean of the top n ChIPs in the population to produce a ratio where:

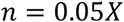

and

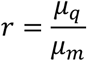

Where μ_q_ is the mean for the top n signals for Q and μ_m_ is the mean of the top n signals for M.

A Score (S) is assigned to each Factor thus:

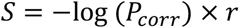

Where P_corr_ is the Benjamini-Hochberg corrected KS 1-tailed p value. Factors are then sorted by Score.

### Implementation of MAGICTRICKS

A tab or comma delimited text file is requested by MAGICTRICKS (lists file). The first column contains a list of all genes of relevance to the experiment (master list). This could be a list of all genes expressed in the system for example, or all genes with a detectable signal. Any number of other columns are then added that contain query lists. For example, a query list may be all genes that go up under some criterion and another may be all genes that go down. The first row is the header and must have unique names for each column.

Using the algorithm above, MAGICTRICKS analyzes each query list and generate a series of output files and sub-directories in the directory containing the lists file. An ‘Accepted_Lists.txt’ file is generated which is the original lists file filtered for genes in the MAGIC Matrix. A Platform_Matrix.txt file is the MAGIC Matrix filtered for genes in the the master list. A series of sub-directories are generated named after each query list in the lists file. Each directory contains sub-directories and files for the stand-alone analysis of that query list. A query_list_summary.pdf contains a bar graph of Factors and Scores with P_corr_ < 0.05 (e.g. Fig. 1b). Query_list_Summary.xls is a spreadsheet containing statistical information for all non-triaged factors.

**Figure 1.**
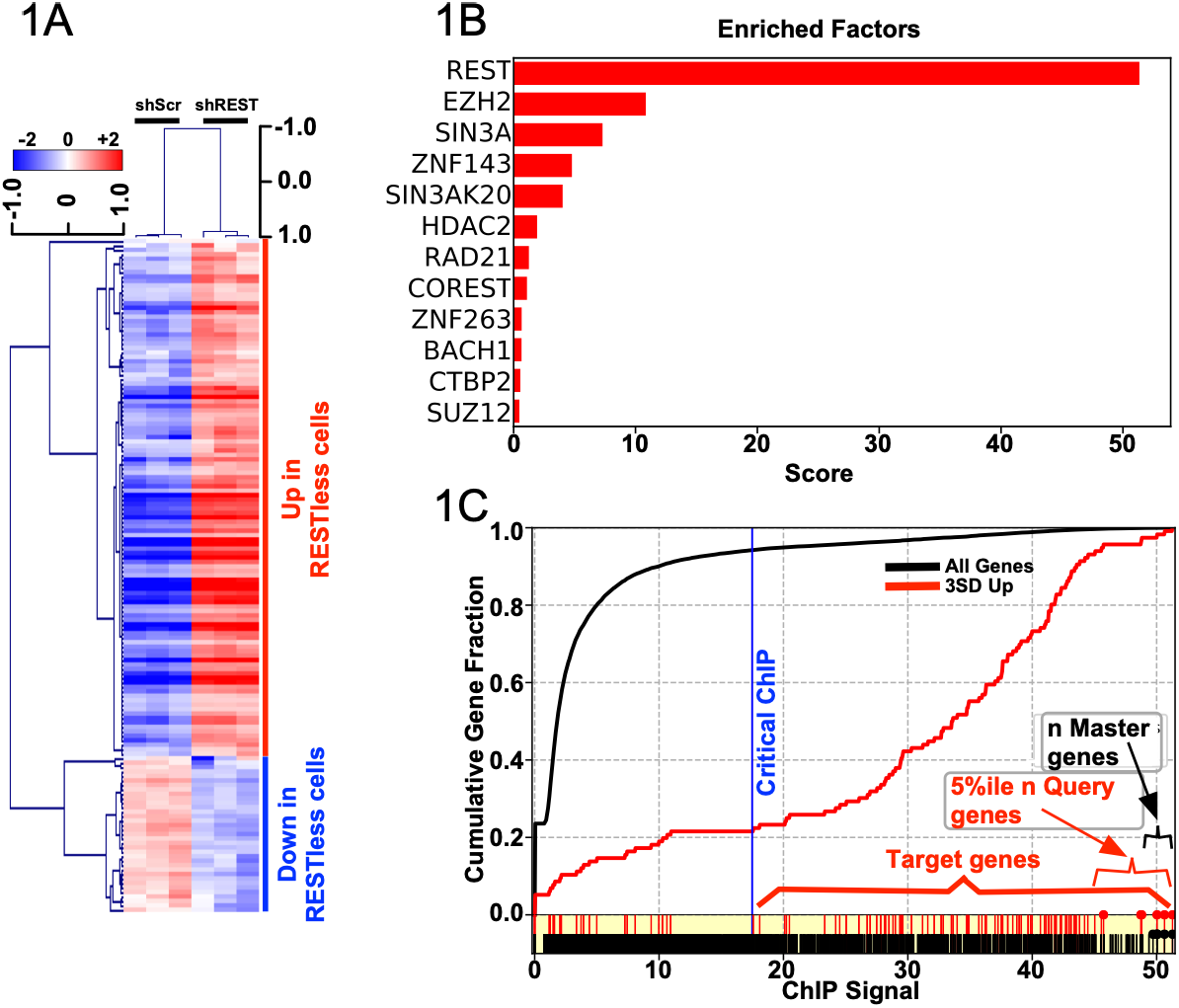
Analysis of MCF7 cells expressing shRNA against REST. **A**) Heatmap showing 119 up-regulated and 35 down-regulated genes by more than 3 standard deviations from the mean fold change at FDR< 0.05. **B**) MAGICTRICKS summary output for up-regulated genes. **C**) Cumulative Distributions Functions of REST ChIP signals extracted from the MAGIC matrix for up-regulated genes (red) or all genes in the analysis (black). Black ticks along the bottom of the graph mark all genes in the analysis. Red ticks mark all up-regulated genes. Lollipops representing the 5%ile of highest ChIP signals in the up-regulated genes list (red) and the same number of highest ChIP signals in the whole analysis (black) are labelled as ‘lollipops’.

Data reported in Query_list_Summary.xls are:

Critical ChIP: ChIP value at d_sup_ (This value is used to determine target genes in the list)
Obs Tail Mean: Average of the 5^th^ percentile ChIP values (n values) in the query list.
Exp Tail Mean: Average of the top n ChIP values in the population.
Tail Enrichment: Ratio of the Obs and Exp Tail Means (r)
Raw P: KS p value
Corrected P (FDR): Benjamini Hochberg corrected p value (P_corr_)
Score: Score (−log(P_corr_)x r)

All Factors associated with P_corr_ < 0.05 are highlighted in bold red.

Query_list_Targets.gmx is a tab delimited file in the GMX format utilized by GSEA^8^. Columns contain all target genes of a Factor whose ChIP signal is greater than the Critical ChIP i.e. the ChIP at which there is the maximal difference between the population and query cumulative. The GMX format will allow this file to be used directly in any subsequent GSEA analysis. The first column contains all genes in the analysis. The second column contains genes in the input list. All subsequent columns contain target genes for the Factor that is named at the top of the column.

A sub-directory called ‘CDFs’ contains graphical displays of the analysis for all factors with P_corr_ < 0.05, named ‘rank’_’factor’.pdf (e.g. 1_NRSF.pdf; rank = 1, factor = NRSF) (Fig. 1c) where ranking is determined by Score. Two cumulative functions are displayed: the black curve is the fractional cumulative of all genes in the master list against the ChIP values, red is the same for query genes. A blue vertical line denotes the Critical ChIP value i.e. ChIP value at d_sup_. Red ticks represent each gene in the query list and black ticks are all genes in the population. Red ticks with circles (‘lollipops’) are those n query genes in the 5^th^ percentile of ChIP values. Black lollipops are genes in the population with the n highest ChIP values.

A second sub-directory called ‘Auxiliary_Files’ is populated with data behind the summary files. The ‘query_list_raw_results.txt’ file contains the same columns as the Query_list_Summary.xls file but has raw data for all factors including those that were triaged and not considered for further statistical analysis. It also contains the KS statistic for each factor. KS statistics with a negative sign denote KS values for triaged factors; the negative sign is used by the algorithm for triage sorting.

Query_list_Sub_Matrix.txt is the MAGIC Matrix filtered for genes in the query list.

Triaged_Factors.txt is a list of factors that were not considered (triaged) because |d_inf_| > |d_sup_|

Triaged_Genes.txt contains all genes in the query that were not in MAGIC Matrix and therefore eliminated from analysis.

A sub-directory named ‘Target_Data’ contains comma separated text files for each Factor with P_corr_ < 0.05. The file contains the list of target genes for that Factor and associated MAGIC Matrix ChIP value. Within ‘Target_Data’ a directory named ‘_FDR_above_5_percent’ contains lists of target genes and associated ChIP values for Factors with an P_corr_ ≥0.05.

### Obtaining MAGICTRICKS

MAGICTRICKS can be downloaded from https://go.wisc.edu/MAGICTRICKS.

## Results

The purpose of MAGICTRICKS is to identify transcription factors and cofactors that are responsible for the major patterns of gene expression changes in a disease situation or other biologic pertubation. To test MAGICTRICKS, we asked whether MAGICTRICKS could predict which transcription factor was knocked down in a cell line. We have shown that REST is a tumor suppressor whose loss drives tumor growth of MCF7 cells in mouse xenograft model of breast cancer. We have previously published on the transcriptome of MCF7 cells infected with lentivirus expressing either a scrambled shRNA or shRNA targeting REST^5,9,10^. This simple system consists of a clonal cell line lacking a single factor. RNA was subjected to hybridization to Nimblegen arrays, and all genes were assigned a Fold Change (FC=log2(shREST/shScramble)) and those with FC>3 Standard Deviations from the mean FC and FDR<0.05 were designated as the ‘Upregulated’ gene list upon loss of REST (118 genes) or ‘Downregulated’ (35 genes not including REST itself). These 153 genes were able to distinguish shScramble from shREST samples in unsupervised heirarchical clustering (Fig.1A). Using 15,445 expressed and 118 upregulated genes (supplemental file 2), MAGICTRICKS predicted REST as the highest scoring Factor associated with upregulated genes in shREST cells (raw p=7.6×10^−55^, FDR = 3×10^−53^, Score = 51.3). MAGICTRICKS also classified the REST corepressors SIN3A, CoREST, HDAC2, and CtBP2 as significant at FDR<0.05.(Fig.1B and supplemental file 3) (see discussion). Figure 1C shows the highly divergent cumulatives of the population of REST ChIP values (black) and those associated with genes induced more than 3 standard deviations of the mean change (red).

For comparison, we used the 3SD gene list in oPossum. Although oPossum did predict REST as the most likely driver (z score = 83) (supplemental file 4), oPossum predicted 45 other transcription factors at z>1.65. Most of the factors have no known physical or functional association with REST. REST is a unique TF in that it has an extended binding motif of 23bp, which likely aided its identification by oPossum. Regardless, oPossum was not able to call any REST cofactors as drivers of the gene list.

The same list yielded REST as the top hit in 13 ENCODE cell lines using Enrichr followed by EZH2 and HDAC2. No other Factors were significant at FDR<0.05 (data not shown)

Next we tested MAGICTRICKS on The Cancer Genome Atlas (TCGA) Nature 2012^11^ provisional breast cancer dataset. We extracted 364 Luminal and 79 Basal-like tumors. Luminal tumors are a subset of breast cancers defined by their robust expression of estrogen receptor alpha (ERα) and associated pioneer factors. They also express the progesterone and her2 receptors. Basal-like tumors are estrogen receptor negative and also lack progesterone and her2 receptors. This is a complex dataset with over 400 samples and heterogeneous tissue. We took all genes up-regulated more than 3SD from the mean fold change in Luminal tumors relative to Basal-like (203 genes) as the gene list for input into MAGICTRICKS and 17814 expressed genes (supplemental file 5). MAGICTRICKS called GATA3, ERα (ESR1) and FOXA1 as the highest scoring factors (GATA3: p=8.4×10^−14^, FDR = 2.9×10^−12^, Score = 7.6; ESR1: p=1.9×10^−11^, FDR = 3.4×10^−10^, Score = 6.8; FOXA1: p=5.7×10^−7^, FDR = 6.7×10^−6^, Score = 3.9). These 3 Factors characterize the luminal breast cancer phenotype and can regulate their mutual expression^12–19^. Importantly, MAGICTRICKS also called ERα cofactors EZH2^20^, P300^21^ and ZNF217^22^ as significant Factors at FDR<0.05 (Fig.2 and supplemental file 6). The same gene list yielded 46 TFs with oPossum at z <1.65 with ERα ranked at 41 (z=2.633). FOXA2 (z=23.2) and FOXA1 (z=20.1) were the top hits (supplemental file 7). Thus, whereas MAGICTRICKS predicted ERα as a principal factor and identified its cohort of pioneer factors and cofactors in ER positive breast cancer, oPossum was unable to clearly underscore the estrogen receptor biology in this test.

**Figure 2.**
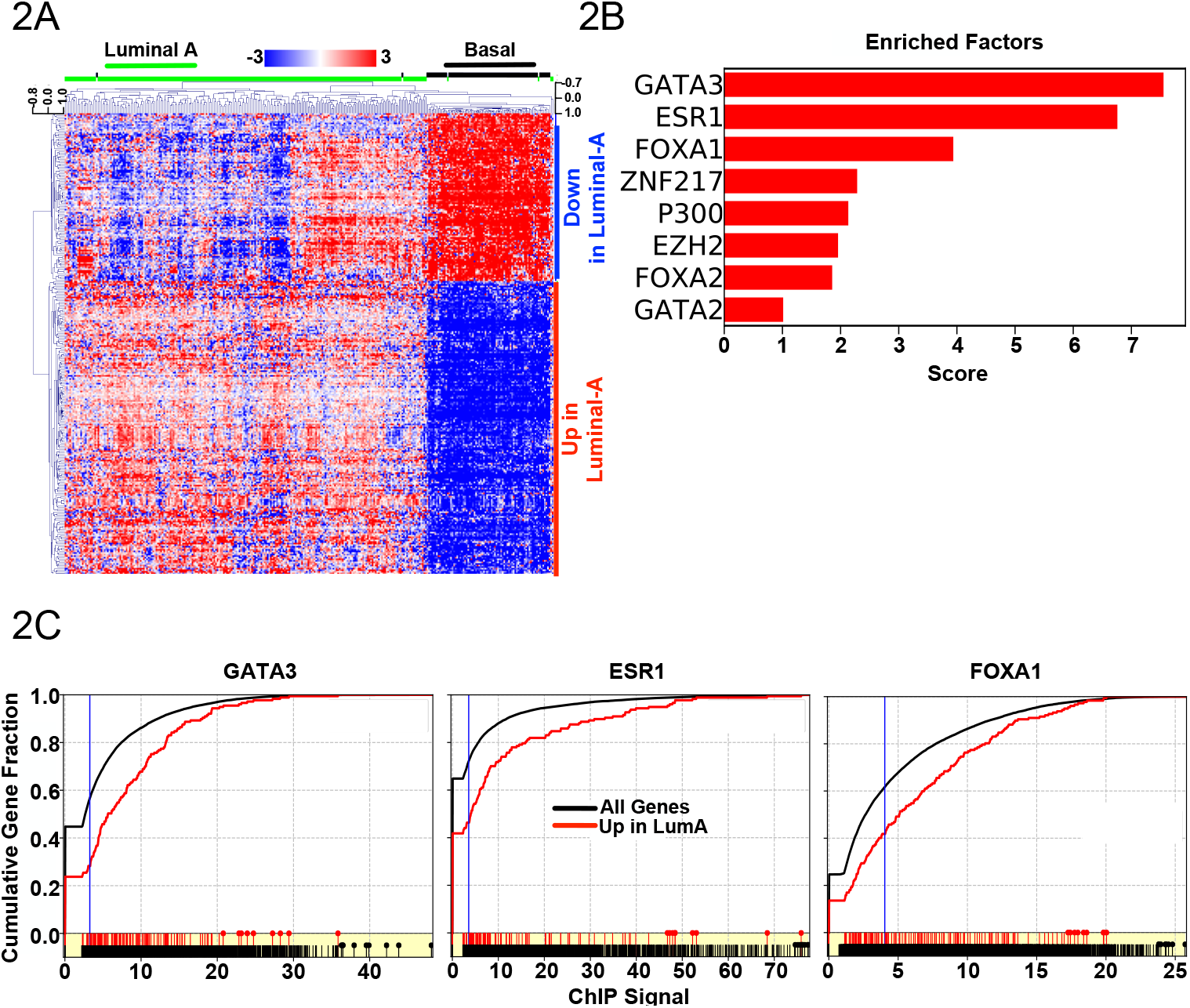
Analysis of Luminal versus Basal-like tumors in TCGA. **A**) Heatmap showing 179 up-regulated and 69 down-regulated genes by more than 3 standard deviations from the mean fold change at FDR < 0.05 across 364 Luminal (232 luminal-A and 132 Luminal-B) and 79 Basal-like tumors **B**) MAGICTRICKS summary output for up-regulated genes. Cumulative Distributions Functions of REST ChIP signals extracted from the MAGIC matrix for up-regulated genes (red) and 14840 background genes in the analysis (black).

At the same FDR cutoff (<0.05), Enrichr yielded ESR1, GATA3, P300, TCF12, EZH2 and FOXM1 (supplemental file 7 – Enrichr tab). Thus there was greater congruance between MAGICTRICKS and Enrichr. Interestingly, wheras Enrichr provided a list of 20 ESR1 target genes, MAGICTRICKS offered 95 Estrogen Receptor targets as defined by having a ChIP signal greater than the Critical Chip (supplemental file 6 – ESR1 targets tab). To test whether MAGICTRICKS was providing a biologically meaningful expanded target gene list rather than merely a large list of genes unrelated to estrogen receptor biology, we performed ontological analysis on target genes from Enrichr and MAGICTRICKS. The 20 ESR1 Enrichr and 95 ESR1 MAGICTRICKS targets were entered into STRING^23–25^ with the 17814 expressed genes as background. Both lists highlighted ‘Estrogen-dependent gene expression’ as the top Reactome^26^ term. Three of the 20 Enrichr genes were associated with ‘Estrogen-dependent gene expression’ at FDR=0.0112 wheras 7 of the 95 MAGICTRICKS genes associated at FDR=5×10^−4^ (supplemental file 8). Importantly, whereas MAGICTRICKS tagged ESR1, FOXA1 and PGR1 as ESR1 targets consistent with known estrogen receptor regulation^27–29^, Enrichr failed to call these crucial genes as targets. Further, under the STRING ‘Biological Processes’ output, the MAGICTRICK target list was tagged with the terms ‘gland morphogenesis’ (FDR=8.5×10-5), ‘branching involved in mammary gland duct morphogenesis’ and ‘mammary gland alveolus development’ (FDR=4.5×10-3) amongst other terms associated with mammary gland development and pregnancy associated remodelling (supplemental file 8). The Enrichr target list was not associated with these terms.

MAGICTRICKS was developed using ChIPseq data derived from immortalized or transformed human cancer cell lines and the above two examples utilize either an immortalized cell line or cancer tissue. To test whether MAGICTRICKS could be used on data derived from non-cancerous, highly heterogeneous, rodent samples, we turned to non-malignant mouse brain tissue. CTCF is a TF that organizes long distance interactions in the genome. Sams et al demonstrated that elimination of CTCF in excitatory neurons (while sparing its expression in other neuron subtypes, astocytes and glia) results in defects in learning, memory and neuronal plasticity^30^. RNA from whole hippocampus (i.e. all cell types in the formation) from WT and CTCF knock-out mice (n=3 per condition) was subjected to RNAseq and the transcriptome data was archived as GSE84175. We generated two gene lists from this dataset: genes up or down-regulated upon CTCF loss at least 1.2x with FDR<0.25 and raw p<0.05 (696 and 432 genes respectively – supplemental file 10) (Fig. 3A). Figure 3B and supplemental file 11 (‘Down’ tab) shows that the 3 highest scoring Factors targeting genes that are down-regulated in CTCF-KO hippocampii are CTCFL, ZNF143 and CTCF itself. Interestingly, for the list of up-regulated genes, MAGICTRICKS yielded MAFF and MAFK as top hits (Fig. 3C and supplemental file 11 ‘Up’ tab) which are associated with Locus Control Regions that require CTCF function^31^.

**Figure 3.**
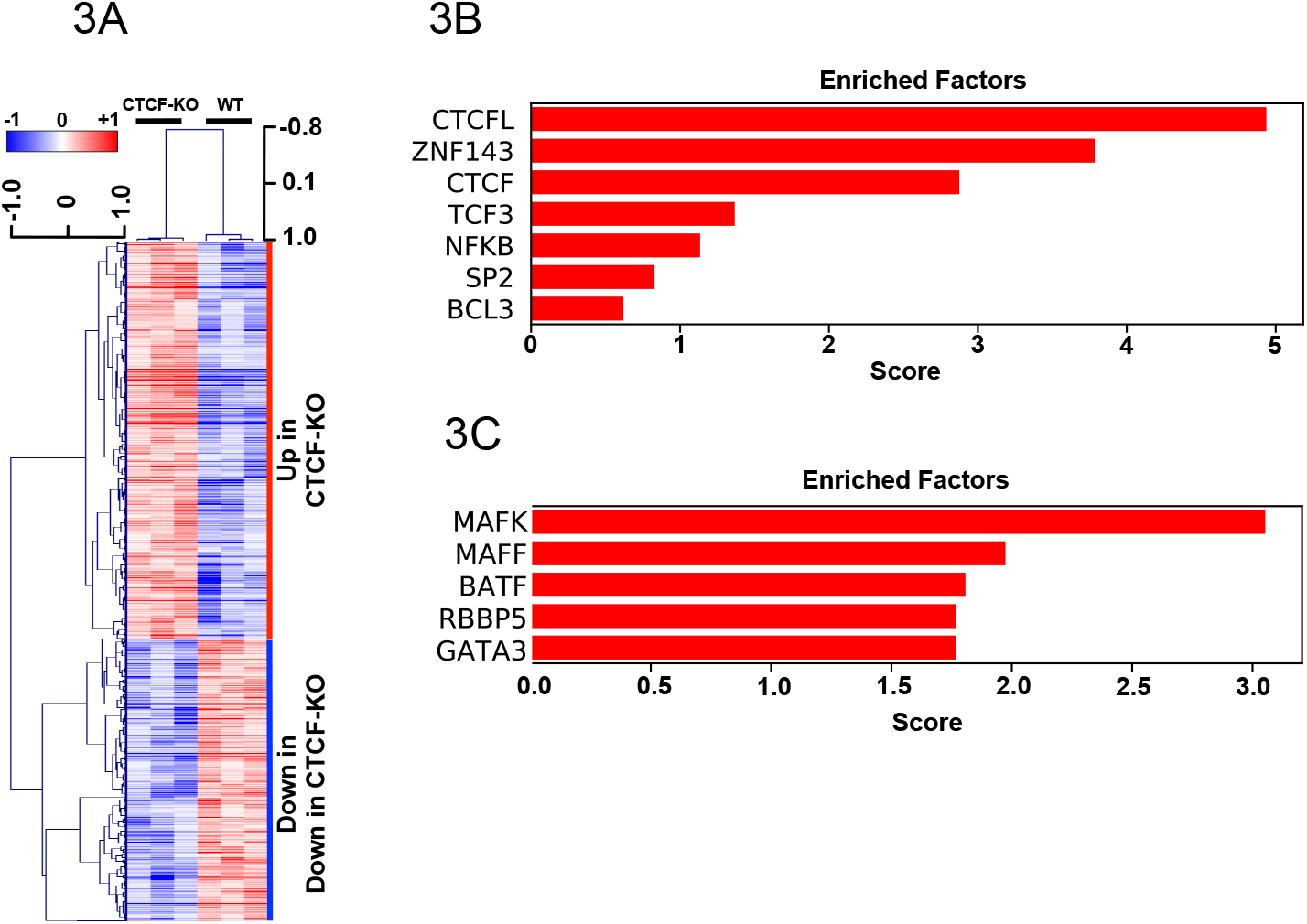
Analysis of WT and CTCF Knock-out hippocampal tissue (GSE84175). **A**) Heatmap showing 696 up-regulated and 432 down-regulated genes changes at least 1.2x with FDR<0.25 and raw p<0.05 **B**) MAGICTRICKS summary output for up-regulated genes. **C**) MAGICTRICKS summary output for genes up-regulated at least 1.2x.

oPOSSUM yielded ZNF354C as the top hit with CTCF ranked number 36 (z=3.8) for down-regulated genes (supplemental file 12 – oPossum tab). Enrichr yielded CTCF as the top hit and was thus more congruent with MAGICTRICKS (supplemental file 1 – Enrichr tab).

In summary, MAGICTRICKS was able to predict which TFs and cofactors drive gene changes in three different types of transcriptomic analysis: a cell line with or without REST, a tumor cohort with or without ERα and brain samples with or without CTCF.

## Discussion

We introduce MAGICTRICKS as a standalone application that predicts transcription factors and cofactors that drive gene expression changes in transcriptomic experiments. In comparison to a commonly used resource, oPossum, MAGICTRICKS predicted factors that were directly relevant to the biology behind the experiment. MAGICTRICKS predicted both the transcription factors whose binding are enriched in the input list as well as cofactors that are consistent with the known biology of the cognate transcription factor.

MAGICTRICKS was developed in response to an unmet need that would allow allocation of transcription factors and cofactors to lists of genes obtained through whole transcriptome experiments. A widely used approach - exemplified by oPossum - rely on searching for short transcription factor binding motifs in promoter proximal sequences. Using hypergeometric tests provides a likelihood of enrichment for the factor. This approach is limited due to the small size of binding sites for most factors; a random 6bp sequence will be found approximately 750,000 times in the human genome. This approach also cannot predict cofactors as cofactors do not bind DNA directly.

An alternative approach relies on comparing a query list of genes with gene lists attributed to factors based on an arbitrary minimal signal or an arbitrary rank cutoff based on ChIPseq experiments. This approach is exemplified by Enrichr ^2^ that accepts input lists and predicts factors based on Chea and ENCODE as well as literature mining. Enricher works by taking the top 2000 ChIPed genes for each Factor in each cell line in ENCODE as the ‘target list’ for that Factor. Fisher tests are then performed to assess association between the query list and the 2000 targets. Enrichr was accurately able to predict REST, ESR1 and CTCF in the 3 examples used in this manuscript. The noticable difference between MAGICTRICKS and Enrichr was in the identification of Factor target genes. Enrichr identifies target genes in the user list by Venn analysis of the user list and pre-defined targets of a Factor; MAGICTRICKS identifies targets by searching for ChIP signals greater than the argument of the Kolmogorov-Smirnov statistic. In the case of ERa when comparing the transcriptomes of Luminal-A and basal-like tumors, ERa targets defined by Enrichr did not include ERa itself, nor PGR or FOXA1 whereas these three key ERa targets^27–29^ were in the set of ERa MAGICTRICKS targets. The larger list of ERa targets defined by MAGICTRICKS were enriched for genes associated with estrogen receptor biology as assessed by ontology (Supplemental File 8).

Though Enrichr is a very powerful, userfriendly and informative resource, the arbitrary boundary of target/non-target genes set at 2000 genes does not take into account the fact that many factors have highly skewed binding profiles: many genes will show some, low level of binding/chip signal for a given Factor. For example, extracting all REST ChIPseq tracks from wgEncodeRegTfbsClusteredV2.bed from the UCSC genome browser (at http://hgdownload.soe.ucsc.edu/goldenPath/hg19/encodeDCC/wgEncodeRegTfbsClustered/) showed 27,386 binding sites across the human genome with 17,971 genes showing a detectable signal within 5Kb of the promoter (data not shown). This thwarts unbiased attempts to define genes as ‘targets’ or ‘non-targets’. MAGICTRICKS overcomes these shortfalls by utilizing complete ChIPseq profiles without parsing Factors into ‘bound’ and ‘not bound’ classes.

Testing MAGICTRICKS on cells transfected with REST shRNA highlights the ability of the algorithm to not only identify transcription factors but also their associated cofactors. MAGICTRICKS called REST as the highest scoring factor but also called SIN3A, HDAC2, CoREST and CtBP2, which are well-characterized REST cofactors^32–35^. EZH2 and SUZ12 were also called as factors that preferentially bind up-regulated genes. Though there are reports of REST interacting with Polycomb^36^, it is likely that Polycomb targets many of the same genes as REST but does so independently^37^. In either case, using a well defined clonal cell line, MAGICTRICKS calls Factors that are either cofactors for REST or co-regulate the same genes. Using gene lists derived from over 300 breast tumor transcriptomes, MAGICTRICKS also correctly called ERα as the principal driver of gene induction in ER positive (Luminal-A) versus negative (Basal-like) tumors. ERα binds chromatin with pioneer factors and MAGICTRICKS sucessfully called GATA2, GATA3, FOXA1 and FOXA2^19^. P300 and EZH2 are ERα cofactors and were called by MAGICTRICKS^20,21^.

In comparison, oPossum was able to identify REST as the principal factor in the first analysis by using Fisher tests of the input list against lists of genes defined as transcription factor targets by proximity to sequence motifs. As expected, oPossum was unable to call any cofactors due to an absence of binding motifs for this class of nuclear protein. With the breast cancer samples, oPossum ranked ERα 41 out of 116 candidate factors. Thus, using oPossum, a naïve researcher studying transcriptomes from Luminal-A and Basal-like tumors would have had difficulty identifying estrogen receptor as a principal factor in the biology of this disease. However, the ERα pioneer factors FOXA1 and FOXA2 were ranked #1 and 2 by oPossum, which would hint at a role for steroid receptors in the disease. Nevertheless, MAGICTRICKS highlighted estrogen receptor biology as a primary driver of transcriptional differences between the 2 groups of tumors.

ENCODE hosts ChipSeq tracks derived from transformed or immortal cell lines and human embryonic stem cells. The MAGIC Matrix incorporates the highest ChIP signal observed across all cell lines within a gene domain for a given factor. This approach was chosen (rather than taking the mean signal for example) because MAGICTRICKS is a discovery platform and taking the maximum value is an attempt to maximize the chance that a particular site may be bound in other systems. A particular concern was that generating a matrix from homogenous cell lines may preclude MAGICTRICKS’s use in analysis of transcriptome data derived from in vivo samples that are not immortalized and likely heterogeneous. However, testing of MAGICTRICKS on GSE84175 demonstrates that RNAseq data derived from control and experimental whole brain extract is handled appropriately. In GSE84175, CTCF was deleted in a subset of post-mitotic cells (eliminated from excitatory neurons but retained in all other neuronal subtypes) ^30,38^. RNA was extracted from whole hippocampus, which included glia and astrocytes. MAGICTRICKS correctly called CTCF and CTCFL as the factors most likely to drive gene changes in the experiment. Interestingly, ZNF143 was also in the top 3 hits. This is in keeping with the recent findings of Mourad and Cuvier^38^ showing that CTCF and ZNF143 coordinate chromatin border domain formation. Thus, MAGICTRICKS was able to call CTCF as well as a known associated factor when provided transcriptome data from a highly heterogeneous tissue where only a small subset of cells were altered in the experiment.

A clear limitation of the current input matrix is that only factors with peaks within 5Kb of the gene body were assigned to a gene: it is clear that a more complete method will take into account distal enhancers that have major roles in global gene regulation. However, despite the absence of distal enhancer information in the MAGIC Matrix, the above three examples show the utility of MAGICTRICKS and MAGIC Matrix using only gene-proximal factor binding. Another limitation is the reliance on a matrix comprised of 161 ENCODE ChIPseq tracks despite there being over 1000 proteins in the human genome that reside in the nucleus. However, the algorithm can accept user defined ChIP seq data and so can evolve as more tracks become available. Users can add their own ChIPseq data to the matrix. To do so, the user would find all ChIPseq peaks within 5Kb of all gene bodies in the MAGIC Matrix and pick the highest to assign to that gene. The sum of all peaks so defined would be normalized to sum to 100,000 and added to the MAGIC matrix as a column.

In summary, MAGICTRICKS is an application for identifying transcription factors and cofactors that preferentially bind lists of genes and provides comprehensive lists of target genes per factor.

## Acknowledgements

I would like to thank Ray Dingledine, Emory University for critical insights into the statistics behind MAGICTRICKS. I thank to Caroline Alexander, John Svaren and Steve Goldstein (University of Wisconsin at Madison) for helpful edits of the manuscript. I would like to thank Queen because ‘It’s a Kind of Magic’.

